# Snow flies self-amputate freezing limbs to sustain behavior at sub-zero temperatures

**DOI:** 10.1101/2023.05.29.541388

**Authors:** Dominic Golding, Katie Rupp, Anne Sustar, Brandon Pratt, John C. Tuthill

## Abstract

All living things are profoundly affected by temperature. In spite of the thermodynamic constraints on biology, some animals have evolved to live and move in extremely cold environments. Here, we investigate behavioral mechanisms of cold tolerance in the snow fly (*Chionea* spp.), a flightless crane fly that is active throughout the winter in boreal and alpine environments of the northern hemisphere. Using thermal imaging, we show that adult snow flies maintain the ability to walk down to an average body temperature of -7 °C. At this supercooling limit, ice crystallization occurs within the snow fly’s hemolymph and rapidly spreads throughout the body, resulting in death. However, we discovered that snow flies frequently survive freezing by rapidly amputating legs before ice crystallization can spread to their vital organs. Self-amputation of freezing limbs is a last-ditch tactic to prolong survival in frigid conditions that few animals can endure. Understanding the extreme physiology and behavior of snow insects is urgently important, as the alpine ecosystems they inhabit are being disproportionately altered by anthropogenic climate change.

## Introduction

Cold temperatures are a significant barrier to animal life in many regions on Earth. Animals that live in polar, boreal, or alpine environments possess physiological and behavioral adaptations to survive and move in extreme cold (Tattersall et al. 2012). Endotherms, such as birds and mammals, generate and conserve body heat to maintain a consistent internal body temperature, even in extreme winter conditions (Blix 2016). However, this strategy is energetically expensive. Perhaps for this reason, most animals are ectothermic — their body heat is derived from the environment. Most ectothermic animals have adapted their physiology and behavior to survive across a range of temperatures. Some insect species migrate before winter begins to avoid decreasing temperatures, while others overwinter in a state of programmed quiescence called diapause (Bale and Hayward 2010; Koštál et al. 2017; Numata and Shintani 2023). These adaptations are necessary as most insects become paralyzed at temperatures below the melting point of water (0 °C). Cold paralysis occurs due to an inability to maintain the membrane potential required for neuromuscular function, a phenomenon called spreading depolarization (Mellanby 1939; MacMillan and Sinclair 2011; Overgaard and MacMillan 2017). Insects in cold environments must also contend with internal freezing of the hemolymph, which is often fatal. The threat of paralysis and death by freezing limits the capacity of most insects to remain mobile and survive at sub-zero temperatures.

The snow fly, *Chionea* spp., is one of the few insects that remains behaviorally active at sub-zero temperatures (**Fig. 1A**). Adult snow flies are found throughout boreal and alpine regions of the northern hemisphere (Byers 1983). In the Pacific Northwest (USA), they are active on the snow from October to May. Snow flies belong to the crane fly family, Tipulidae, and resemble other crane flies except for a complete absence of wings and flight musculature. Female snow flies take advantage of the increase in thoracic space for additional egg storage (Byers 1969).

**Figure 1.**
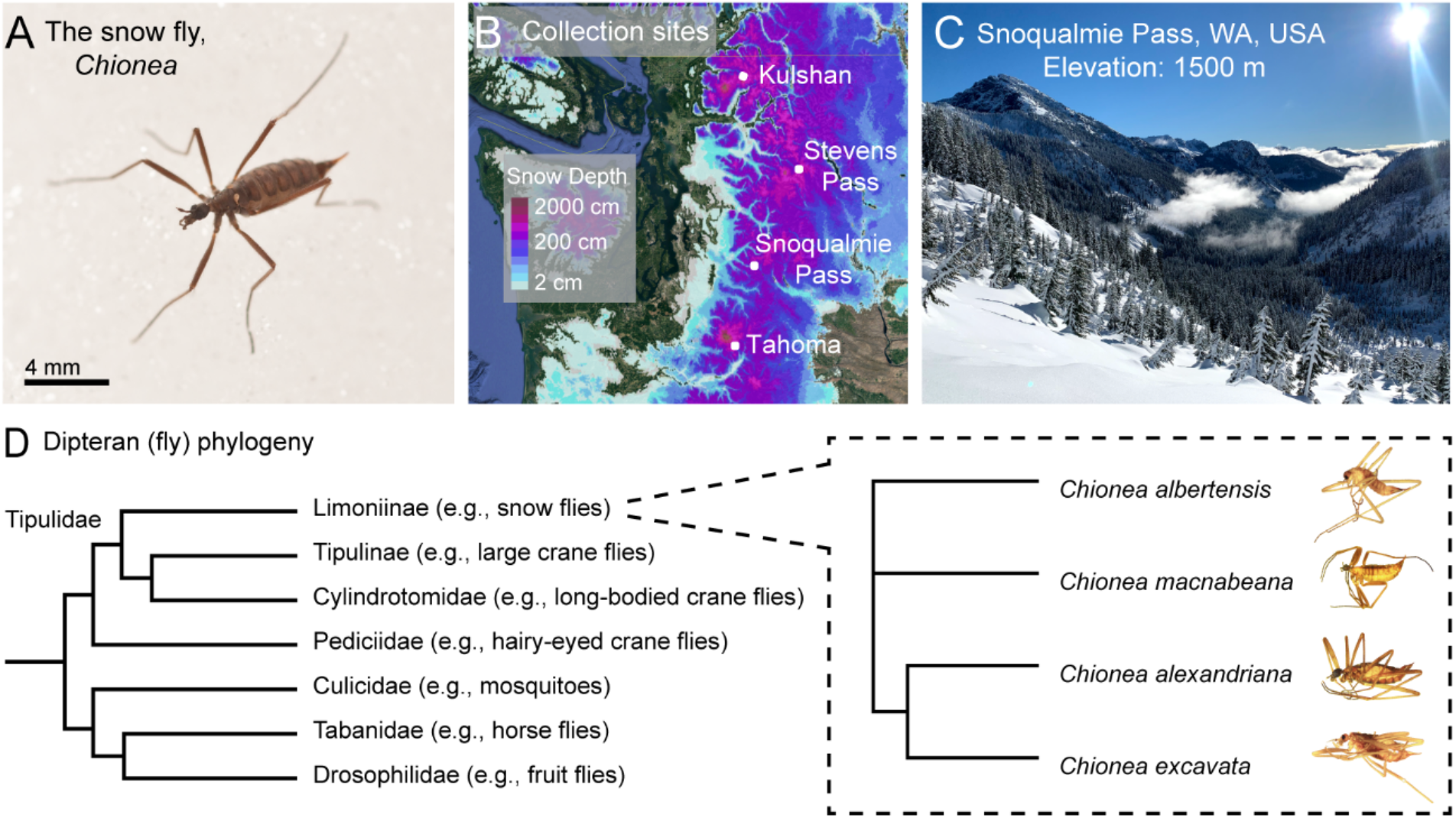
The North American snow fly, *Chionea*. (**A**) Lateral view of a female *Chionea alexandriana*. Photo © Bryan Kelly-McArthur. (**B**) Collection locations of snow fly specimens used for cold tolerance experiments. The four species used in this study significantly overlapped in distribution across collection sites. Collections of flies used for experiments took place from December of 2020 to November of 2022. Flies were found from late October through early May each year. Snow depth is shown for a typical collection day (Feb. 25, 2022). (**C**) Our most popular snow fly collection site, which is representative of the alpine and boreal environments in which snow flies are active throughout the winter. (**D**) Phylogeny of snow flies used in this study. Branch lengths do not correspond to the length of evolutionary time between nodes. Snow flies are a genus within Tipulidae, commonly referred to as the crane fly family. Crane flies are one of the largest and oldest families within the Dipteran order, with over 15,270 species identified (de Jong et al. 2008).

The life cycle of snow flies is poorly understood. Byers (1983) speculated that they deposit eggs in the subnivean zone, beneath the snowpack, in tree litter or moist soil, and that the larvae consume plant detritus or rodent feces during the summer, pupating during late fall or early winter. Adult snow flies are not typically found in the summer, as they have been shown to prefer temperatures around -3 °C (Wojtusiak 1950). We have observed snow flies running on the snow at ambient temperatures as low as -10 °C. They have a significantly longer lifespan than other crane flies, living as long as two months (Byers 1983). Snow flies are not known to eat as adults (Byers 1983), which is not unusual for a crane fly species. Rather, it seems that their primary reason for being active on snow is to locate a mate. Two potential advantages of searching for mates in winter are the uniformity of the snow as a substrate and the absence of predators.

The thermal limits and dynamics of snow fly behavior have not previously been characterized, probably because they are challenging to collect from the wild and cannot be bred in the lab. To overcome these obstacles, we created a crowd-sourced science project (www.snowflyproject.org) that leveraged the expertise of backcountry skiers and mountaineers to collect snow flies from remote alpine regions of the Pacific Northwest. During summer months, we also collected other species of winged crane flies in Western Washington for comparison. We then used thermal imaging to measure their temperature and behavior when subjected to sub-zero temperatures in the lab.

We found that snow flies possess a remarkable ability to sustain locomotion at temperatures well below the paralysis threshold of other crane flies. Our results suggest that snow flies possess adaptations that allow their neurons and muscles to overcome thermodynamic constraints on membrane potential maintenance, ion channel gating, and action potential conduction (Hodgkin and Huxley 1952; Schwarz 1979). We also discovered that snow flies can self-amputate freezing limbs, which prevents the fatal propagation of ice to other parts of the body. Limb self-amputation is common in crane flies (Needham 1953), but is typically triggered by mechanical stimuli, such as during predation (Maginnis 2006). We propose that snow flies possess a unique capacity to rapidly amputate legs using thermosensory detection of freezing body fluids.

## Results

From 2020-2022, we and volunteers collected 256 adult snow flies, primarily from alpine regions of Washington (**Fig. 1B)**, as well as from Colorado, Vermont, British Columbia and Yukon. In Washington, we typically found snow flies running on freshly fallen snow at, or close to, the tree line (**Fig. 1C, Video S1**), between elevations of 1200 to 2000 m, and at an average daily temperature of 0 °C. Collected snow flies were individually housed at 1 °C. We conducted cold tolerance experiments on 39 males and 38 females from four species collected in Washington (**Fig. 1D**): *Chionea alexandriana* (55), *Chionea excavata* (14), *Chionea albertensis* (4), and *Chionea macnabeana* (4). We determined the species of each snow fly using a combination of morphological identification (Byers 1983) and DNA barcoding (Hebert et al. 2003). We did not observe differences in the behavior of these four species, so we pooled data for further analysis.

We first sought to establish the minimum temperature at which snow flies can sustain behavior. We were unable to use the traditional method of measuring the temperature of the flies with a thermocouple (Sinclair et al. 2015), because it prevented the flies from moving. Instead, we used infrared imaging with a thermal camera (Palmer et al. 2004; Tattersall 2016) to record the cuticular temperature and behavior of individual snow flies on a cold plate (**Fig. 2A**). We validated our thermal camera measurements with a thermocouple (**Fig. S1**). During each experiment, the temperature of the cold plate decreased by 0.8 °C/min over 25 minutes (**Fig. 2B**). We manually tracked snow fly movement in the thermal imaging video, from which we extracted the insect’s walking velocity and temperature (**Fig. 2B-C**). The average rate of snow fly cooling was 0.5 °C/min and the temperature of the snow fly was typically a few degrees warmer than the cold plate. We attribute this offset to the thermal gradient between the cold plate and surrounding air, which was 5 °C (see **Methods**). We also observed a thermal gradient between the surface of the snow and the snow fly in the wild (**Video S2**). Therefore, snow flies do not appear to generate substantial heat through muscle contraction, as, for example, bees do (Heinrich 1980).

**Figure 2.**
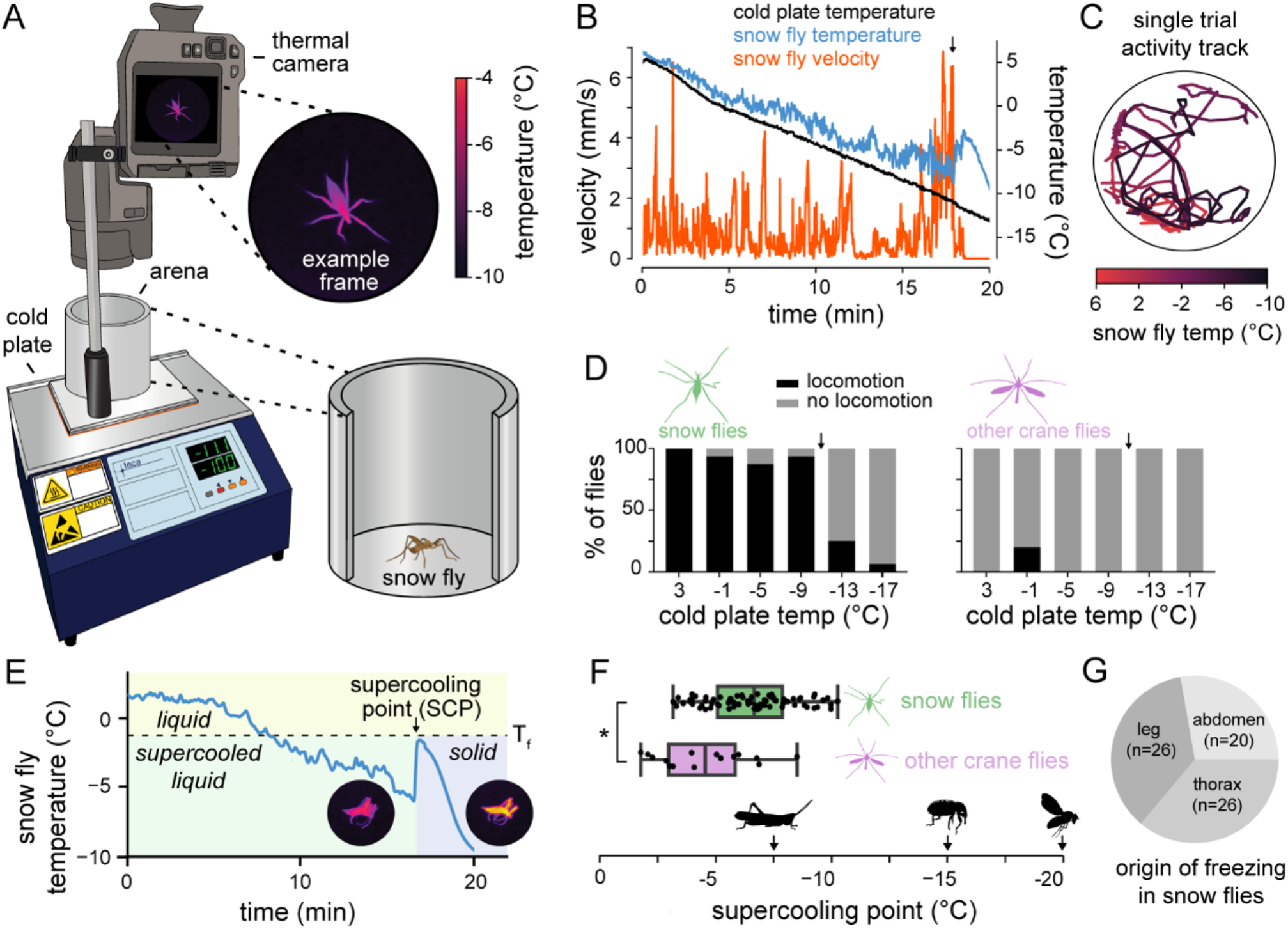
Snow flies sustain locomotor behavior at sub-zero temperatures until they freeze. (**A**) Schematic of the thermal imaging setup used to record snow fly activity and temperature. (**B**) The cuticular temperature of the snow fly (blue), cold plate temperature (black), and fly velocity (orange) during the first 20 minutes of one experimental trial. The fly sustains consistent motor coordination until 18 minutes in, when a rapid temperature increase indicates that the fly has frozen (black arrow). Afterward, all movement ceases. (**C**) Position and temperature of the snow fly within the arena over the course of the trial shown in **B**. (**D**) Snow flies sustain locomotion at sub-zero temperatures (n = 16 trials), while other crane flies do not (n = 5 trials). For each cold tolerance trial (e.g., Panel **B**), we manually scored fly locomotion within 6 different windows (cold plate temperatures of 3, -1, -9, -13, -17 +/- 0.5 °C). Black arrows indicate the average cold plate temperatures at which snow flies and other crane flies froze. (**E**) A second example trial, illustrating that below the freezing point (T_f_), snow flies are supercooled. At the supercooling point (SCP), ice nucleation occurs within the snow fly and ice spreads throughout the body. This is indicated by a rapid increase in the fly’s temperature. Note that this temperature trace was smoothed with a Gaussian filter for presentation, while the trace in **B** was not. (**F**) Distribution of SCPs across snow flies (green; n = 73 trials) and other crane flies (purple; n = 16 trials). The SCP of snow flies is lower than that of other crane flies (two sample t-test, \**p* <0.001). The SCP of short-horned grasshoppers (*Stenocatantops splendens*; Zhu et al. 2013), alfalfa weevils (*Hypera postica*; Armbrust et al. 1969), and fruit flies (*Drosophila melanogaster*; Andersen et al. 2015) are indicated below for comparison. (**G**) Ice nucleation typically occurred in the snow fly’s abdomen, legs, or thorax.

Snow flies continued to move at cold plate temperatures well below 0 °C, with most sustaining locomotion even when the cold plate dropped to -9 °C (**Fig. 2D**). For comparison, we conducted equivalent experiments on other crane flies that we collected locally from Western Washington during summer months (see **Methods**). We found that summer-active crane flies were generally unable to sustain locomotion at cold plate temperatures below 0 °C (**Fig. 2D, Video S3**). Soon after being placed on the cold plate, they entered a coma-like state in which their movement was minimal and uncoordinated. Thus, snow flies appear to possess a unique ability to remain behaviorally active at sub-zero temperatures that incapacitate other insects, including their relatives in the crane fly family.

We observed that the point at which snow flies ceased movement was preceded by a sudden increase in body temperature (**Fig. 2E, Video S4**). This sudden warming is an indication of ice formation within the snow fly’s hemolymph (Sinclair et al. 2015). Freezing is an exothermic process, releasing heat as the liquid molecules rearrange into a crystalline molecular structure (Duman 2001). Internal ice formation was almost always lethal; only 3/77 snow flies survived after freezing occurred within their body. This low survival rate indicates that *Chionea* spp. are generally not freeze tolerant.

Cold-tolerant insects that cannot survive internal ice formation often exhibit physiological and behavioral traits that help them avoid fatal freezing events. For example, some freeze-avoidant insects synthesize sugars and proteins that act as antifreeze (Duman et al. 1995; Duman 2001). This allows them to take advantage of the phenomenon of supercooling, by which small volumes of water can remain in a liquid state to -40 °C or lower (Zachariassen et al. 2004). Supercooling is common in small insects, as their hemolymph primarily consists of water and lacks nucleating particles that are required for the ordered conglomeration of water molecules during ice formation. The temperature at which an insect freezes is referred to as its supercooling point, or SCP (Bale 1987; Sinclair et al. 2015).

We measured the SCP of snow flies (n = 73 trials) and crane flies (n = 16 trials) from the same cold tolerance experiments described above (**Fig. 2F**). Because they spend most of their adult lives exposed to sub-zero temperatures, we expected that the SCP of snow flies would be significantly depressed compared to summer-active crane flies. The SCP of snow flies was lower than that of other crane flies (−6.6 ± 1.9 °C vs -4.6 ± 2.0 °C, *p* < 0.001, two-sample t-test), but it was also close to or higher than other insect species that are not cold tolerant, such as grasshoppers, weevils, and fruit flies (**Fig. 2F**). For example, the SCP of *Drosophila melanogaster* has been found to be -20 °C, although fruit flies are unable to sustain locomotion below 0 °C (Andersen et al. 2015). Thus, snow flies have a unique ability to sustain behavior until the moment they freeze, even though they freeze at similar temperatures as summer-active insects.

We also examined the site of ice formation within the snow fly’s body. Although the posterior region of the abdomen and distal segments of the tarsi were typically the coldest regions of the snow fly, this was not always where ice nucleation occurred: 28% of terminal freezing events originated in the abdomen, 36% in the thorax, and 36% in a leg (**Fig. 2G**).

When we inspected the thermal imaging videos more closely, we noticed that snow flies frequently self-amputated their legs following internal ice formation (**Fig. 3A, Video S5**). Limb self-amputation, also known as autotomy, was relatively common: over half of the snow flies lost at least one leg during our experiments (43/77 flies). Amputation occurred 31% of the time after freezing originated in a leg (60/194 instances). The median number of limbs lost across trials was one, although flies lost as many as five limbs prior to the rest of the body freezing. Limb self-amputation in our experimental setup followed a typical pattern. The snow fly’s leg often became stuck to the plate due to freezing condensation. As the snow fly struggled to free itself, the tip of the leg froze, indicated by a sudden increase in temperature at the tarsus, radiating proximally toward the tibia and femur (**Fig. 3A-B, Video S5**). We quantified the time course of ice propagation within the leg (**Fig. 3B)**. The median time it took for ice to propagate from the snow fly’s tarsus to the tibia was 533 ms (n = 44 instances). During successful amputations, the leg detached from the body before the wave of ice reached the trochanter. Amputation occurred at a median delay of 2.5 seconds following ice formation in the tarsus (n = 60 instances). Failure to detach the leg following ice nucleation in the tarsus resulted in ice formation within the body, which was fatal for the fly (**Video S5**).

**Figure 3.**
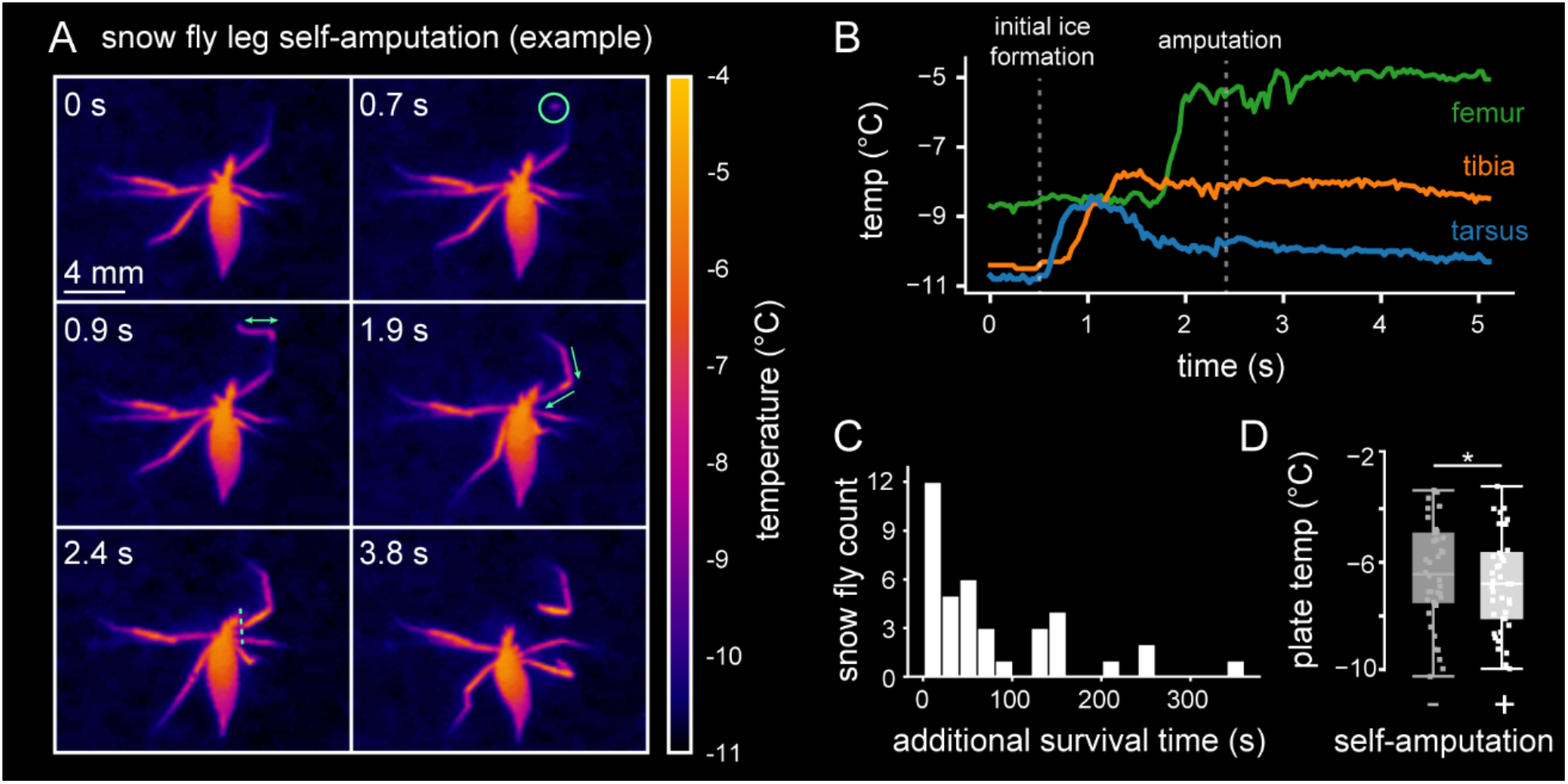
Snow flies evade fatal freezing events by self-amputating limbs (autotomy). (**A**) An example of snow fly leg amputation. At 0 s, the tibia and tarsus of the fly’s right front leg are at the same temperature as the cold plate (−11 °C). At 0.7 s, freezing occurs in the mid tarsus (0.7 s, green circle) and spreads bidirectionally toward the tip of the tarsus and the tibia (0.9s, green arrow). Ice crystallization propagates through the tibia and reaches the femur (1.9 s, green arrows). Amputation at the femur/trochanter joint occurs at 2.4 s, before freezing can propagate through the trochanter and to the body. The fly pulls away from the detached leg while the leg flexes, with the body of the fly remaining unfrozen (3.8 s). (**B**) Time course of freezing in the tarsus (blue), tibia (orange), and femur (green) for the example in **A**. The time of detachment (white dotted line) indicates that autotomy is rapid, occurring within 1.7 s of the initial freezing event in the tarsus. (**C**) Successful amputation increases snow fly survival time in lab experiments (n = 43 trials). The average additional time of survival was 77 s. (**D**) Snow flies that amputate at least one limb (n = 43 flies) survive to a lower cold plate temperature than snow flies that retain all limbs (n = 34 flies) during cold tolerance experiments (two sample t-test, \**p* = 0.01).

Leg amputation also appears to occur frequently in the wild: nearly 20% (15/77) of the snow flies we collected were missing one or more legs. In rare cases, we even observed snow flies effectively navigating complex snowy terrain with only three legs (**Video S6**).

Since amputation often followed leg freezing, we speculated that this could be a mechanism to prolong snow fly survival under extreme environmental conditions. We found that snow flies that amputated a limb following ice nucleation in the tarsus survived an average of 77 additional seconds before freezing in our experimental setup (**Fig. 3C**). Snow flies that amputated limbs (n = 43 flies) froze at significantly lower cold plate temperatures than those that did not (n = 34 flies) (**Fig. 3D**; two-sample t-test, -11.6 ± 1.2 °C vs -10.7 ± 2.0 °C, *p* = 0.01). There was no significant difference in SCP between snow flies that amputated legs and those that did not (two-sample t-test, -6.8 ± 1.8 °C vs -6.2 ± 2.0 °C, *p* = 0.20). Overall, snow flies that amputated frozen legs survived longer and to lower cold plate temperatures.

Crane flies and some other insects can self-amputate legs in response to pulling, such as when they are captured by a predator (Needham 1953; Maginnis 2006). We tested whether leg self-amputation in snow flies could be initiated by a mechanical stimulus applied to the leg (**Fig. 4A**). We held one of the legs captive using forceps and prodded the captive leg with a mechanical probe (**Video S7**). The mechanical stimulus failed to produce leg self-amputation in snow flies. However, two-thirds of these same snow flies later lost at least one limb during cold tolerance experiments (**Fig. 4A**). In contrast, over half of the other crane flies self-amputated a limb when we applied the same mechanical stimulus to their legs. We never observed other crane fly species self-amputate legs during cold tolerance experiments (**Fig. 4A)**.

**Figure 4.**
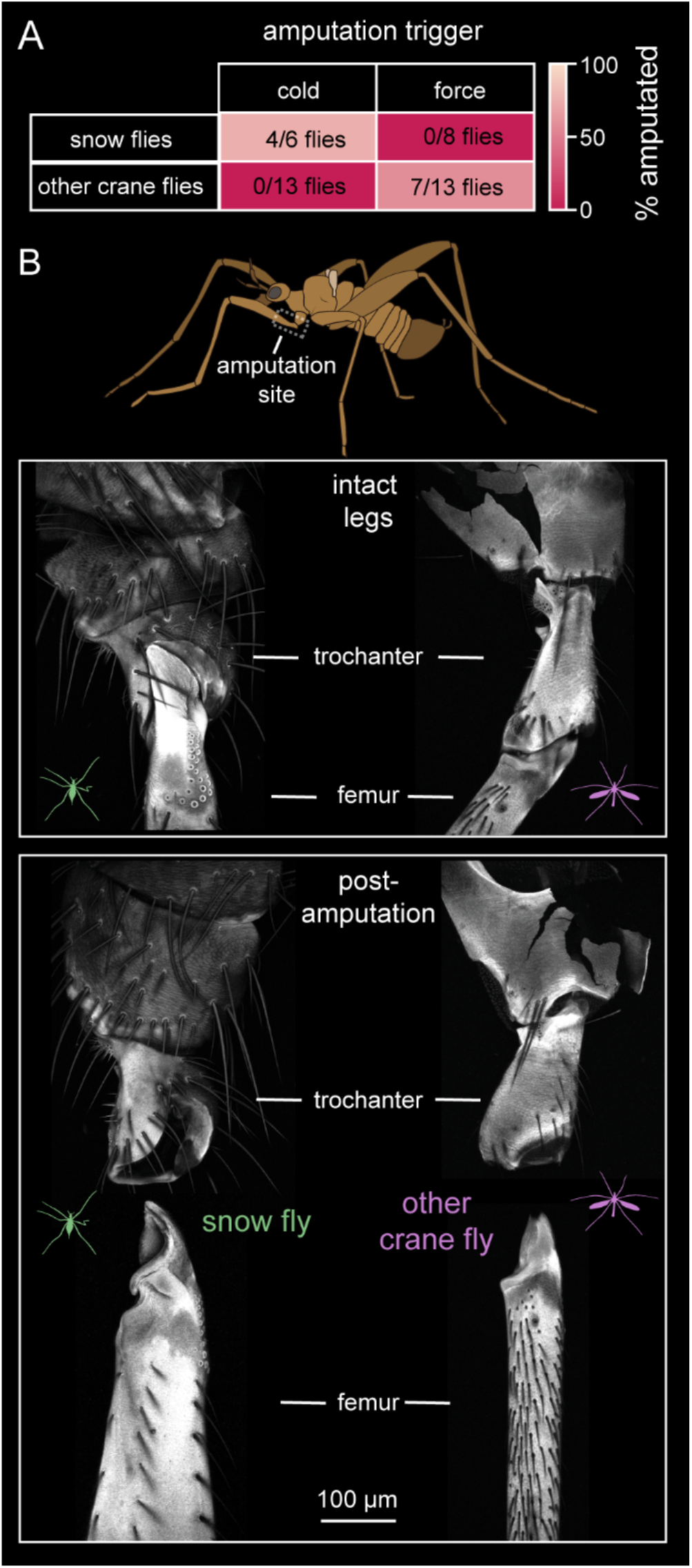
Leg self-amputation occurs at the same site in the leg, but is triggered by distinct stimuli, in snow flies vs. other crane flies. (**A**) Snow flies self-amputate legs in response to freezing, but not pulling with forceps. Other crane flies self-amputate legs in response to pulling, but not freezing. The same flies were used for both mechanical and cold plate autotomy experiments. Two snow flies had died prior to the cold plate autotomy experiments. (**B**) The morphology and amputation site at the femur/trochanter joint are consistent across snow fly (left) and crane fly (right) legs. Images show cuticle autofluorescence (emission wavelength of 633 nm).

Leg amputation in snow flies and other crane flies always occurred at the same point within the leg: the joint between the femur and the trochanter (**Fig. 4B**). The structural similarities across snow flies and other crane flies suggest that amputation is mediated by activation of similar muscles, which have been previously described in other insects (Schindler 1979). We speculate that leg self-amputation originally evolved as an anti-predation mechanism in summer-active crane flies and was subsequently co-opted by snow flies to prevent propagation of internal freezing and prolong survival in extreme cold.

## Discussion

We used thermal imaging to show that snow flies can sustain behavior at body temperatures down to a minimum of -10 °C or mean of -6.6 °C. This capacity seems to defy the thermal constraints on neuromuscular function that have been well-characterized in other insect species (Findsen et al. 2014). Most insects are chill sensitive, meaning that they are paralyzed below a chill coma temperature threshold (Findsen et al. 2014; Robertson et al. 2017). At this chill coma threshold, spreading depolarization of the nervous system leads to loss of neuromuscular coordination (MacMillan and Sinclair 2011; Overgaard and MacMillan 2017; Robertson et al. 2020). For example, the chill coma threshold varies from 8 °C to -2 °C in different *Drosophila* species (Andersen et al. 2015). All other crane flies we tested were chill sensitive, with a chill coma above 0 °C. In contrast, snow flies never lose neuromuscular coordination, but sustain behavior up to the point at which they freeze and die. That snow flies move and respond to external stimuli under these conditions suggests that they possess adaptations that allow their neurons and muscles to overcome thermodynamic constraints on ion pumps, channel gating, synaptic transmission, and muscle contraction.

Below 0 °C, the snow fly body fluids are in a supercooled state. Small volumes of water within insects have been shown to supercool to temperatures as low as -30 °C (Zachariassen et al. 2004). Snow flies may produce antifreeze compounds within their hemolymph to depress the SCP (Duman 2001; Vanin et al. 2008). However, some summer-active insects are capable of supercooling to -8 °C without detectable antifreeze or cryoprotectants (Storey and Storey 1996). Although we determined that the SCP of snow flies is 2 °C lower than summer crane fly species (**Fig. 2F**), this difference suggests that production of antifreeze or cryoprotectants in snow flies is limited. Instead, they likely avoid exposure to temperatures close to their SCP by burrowing into crevices in the snow, where temperatures are consistently close to 0 °C. We also found that snow flies are not able to survive freezing, unlike some species of overwintering insects (Teets et al. 2023).

Although snow flies are not exceptional in their ability to prevent or survive freezing, they have a unique capacity to maintain neuromuscular function at sub-zero temperatures, a trait shared by few other insect species. The only other species that are known to sustain locomotion at similar minimum temperatures are the Alaskan beetle, *Pterostichus* spp. (Baust 1972), the Tasmanian snow scorpionfly, *Apteropanorpa* spp. (Palmer et al., 2004), the Himalayan glacier midge *Diamesa* spp. (Kohshima 1984), and the ice crawler, *Grylloblatta* spp. (Morrissey and Edwards 1979; Schoville et al. 2015). Notably, ice crawlers also live in alpine regions of the Pacific Northwest, where they are the only known predator of snow flies (Pritchard and Scholefield 1978; Schoville et al. 2015).

What are the advantages of maintaining constant motion until the moment they freeze? In the wild, snow flies may improve their chances of surviving cold night-time temperatures if they search for an insulated place to take refuge (*e*.*g*., under the snow). Another major advantage of sustaining locomotion close to their supercooling limit is that it allows snow flies to disperse and reproduce in an environment that is mostly devoid of other animals, including predators. We and others (Cannings and Kelly-McArthur 2021) have observed snow flies audaciously mating in full view on the surface of the snow for thirty minutes or more.

We found that snow flies rapidly self-amputate freezing limbs to prevent ice from spreading to the rest of the body. To our knowledge, amputation has not been previously described as a mechanism to avoid freezing and prolong survival at cold temperatures. Limb self-amputation occurs in many species, including reptiles, amphibians, mammals, birds, fish, echinoderms, crustaceans, spiders, and insects (Maginnis 2006; Fleming et al. 2007). It is typically used to avoid capture by predators, reduce cost of injury to a limb, escape non-predatory entrapment, or survive complications during molting (Eisner and Camazine 1983; Fleming et al. 2007; Emberts et al. 2017; Emberts et al. 2020). In snow flies and other crane flies, self-amputation consistently occurs at the joint between the trochanter and femur (**Fig. 4B**). Specialized muscles that control amputation at this joint have been described in stick insects and crickets (Schindler 1979; Frantsevich and Wang 2009; Zill et al. 2017). Based on similarities in the breakage plane (**Fig. 4B**) and snow fly leg musculature (Byers 1983), snow flies appear to use a similar amputation mechanism.

The key difference between leg amputation in snow flies and other insects appears to be the triggering stimulus. Many insects, including other crane flies, self-amputate legs in response to mechanical stimuli, such as pulling on the leg (Emberts et al. 2019). The receptors responsible for sensing mechanical stimuli and triggering leg amputation in other insects are likely campaniform sensilla (Schindler 1979; Frantsevich and Wang 2009). However, we found that mechanical manipulations never triggered leg amputation in snow flies (**Fig. 4A**). We hypothesize that leg self-amputation in snow flies may instead be triggered by thermosensory neurons that detect the temperature increase following ice crystallization of the hemolymph. The rate of ice propagation (533 ms from the tarsus to the femur) would provide ample time to execute this leg amputation reflex.

We propose that the ability of ancestral crane flies to self-amputate limbs may have predisposed snow flies to adapt to their unique lifestyle of wandering cold, snowy environments. However, these ecosystems are rapidly changing due to anthropogenic climate change. Washington is on track to lose 46% of its end-of-winter snowpack by the 2040s and 70% by the 2080s, compared to the 20th century average (Snover et al. 2013). The decrease in snowfall in the Pacific Northwest and across the planet will likely imperil snow flies and other animals that rely on snow for survival (Marshall et al. 2020). We may have limited time to study these species before they disappear altogether.

## Acknowledgements

We thank Kameron Decker Harris, Meira Lifson, Samantha Kunze Garcia, Sydney Cannings, Robert Hahn, Abby Person, and Andres Torres for assistance with snow fly collection. We thank Sydney Brannoch for creating the schematics in Fig. 2A and 4B. We thank Bryan Kelly-McArthur for use of the *Chionea alexandriana* photograph used in Fig. 1A. We thank Alessia Gallio and Marco Gallio for sharing their protocol for DNA barcoding. We thank Katie Marshall, Bing Brunton, and members of the Tuthill Lab for helpful discussions and comments on the manuscript. The cartoon depictions of the short-horned grasshopper (created by Fernando Campos De Domenico) and fruit fly (created by Ramiro Morales-Hojas) in Fig. 3C were obtained from the Phylopic website (https://www.phylopic.org). The cartoon depiction of the alfalfa weevil was created using a reference image from Jennifer C. Girón Duque, from IPM Images (https://www.ipmimages.org). We thank members of the Tuthill Lab for comments on the manuscript. This work was supported by a Searle Scholar Award, a Klingenstein-Simons Fellowship, a Pew Biomedical Scholar Award, a McKnight Scholar Award, a Sloan Research Fellowship, the New York Stem Cell Foundation, and a UW Innovation Award to JCT. JCT is a New York Stem Cell Foundation – Robertson Investigator.

## Methods

### *Chionea* collection and species identification

256 adult specimens of *Chionea* spp. were collected from October of 2020 to late November of 2022. Location coordinates for each collected snow fly are included in **Table S1**. Generally, specimens were found from 10 am to 4 pm during overcast and sunny days after snow had fallen the previous night. On average, flies were captured at a temperature of 0 °C, based on weather data from the Visual Crossing Weather Query Builder (https://www.visualcrossing.com/weather/weather-data-services). Specimens were collected at an average elevation of 1485 m (minimum of 224 m, maximum of 3428 m). Snow depth data in Figure 1B is from the US National Operational Hydrological Remote Sensing Center (https://www.nohrsc.noaa.gov/earth/). All flies were individually kept within 5 mL snap-cap centrifuge tubes, provided a drop (about 0.05 mL) of maple syrup diluted in water, and housed in a refrigerator at 1 °C. After completing experimental trials, dead snow flies were preserved in 95% ethanol at -20 °C. We examined the morphology of 101 dead flies to identify species as described in Byers, 1983. We subsequently used DNA barcode sequencing to confirm our species identifications by comparing a 640 bp region of cytochrome *c* oxidase I subunit I (COI-5P) from each fly. We extracted genomic DNA from leg tissue (n = 245) using 10.1 µL of 100 mM of Tris-Cl (pH 8.0), 1 mM EDTA, 25 mM NaCl, and freshly added 2 mg/mL proteinase K. Each sample was incubated at 37 °C overnight, followed by 5 minutes at 95 °C to inactivate proteinase K. Extracted DNA was then amplified using PCR. We accomplished this by adding 1 µL of DNA after extraction to a solution of 8.8 µL H_2_O, 2X Phusion Master Mix, and 0.1 µL of forward primer (LCO1490) and 0.1 µL of reverse primer (HCO2198). Amplified DNA was sent to the Genewiz laboratory in Seattle, Washington for Sanger sequencing. Traces were uploaded to the Barcode of Life Data System (BOLDSystems) online database where a taxon ID tree was generated using the MUSCLE alignment algorithm (Edgar 2004). The taxon tree included five branches that correlated with our morphological species identifications: *C. alexandriana, C. albertensis, C. jellisoni, C. excavata, and C. macnabeana*. Of these species, *C. alexandriana* was most common (68% of captured flies), followed by *C. excavata* (18%), *C. albertensis* (4%), *C. jellisoni* (4%), and *C. macnabeana* (3%). Behavioral experiments used a subsample of 77 flies from four species in Washington State collected from December of 2020 to November of 2022 (*C. alexandriana, C. albertensis, C. excavata, and C. macnabeana*; **Fig. 1D**).

### Collection and species identification of other crane flies

Adult crane fly specimens (n = 26) were collected from late May to early September of 2022 in Seattle, Gig Harbor, and Toledo, Washington (**Table S2**). Each fly was individually housed at room temperature (approximately 20 °C) in plastic food storage containers lined with wetted paper towels. The DNA barcode sequence of the crane fly used for confocal imaging in **Fig. 4B** was obtained using the snow fly DNA barcode sequencing protocol described above. Species identification for the crane fly depicted in **Fig. 4B** was determined by using the BOLDsystems identification engine. From the cytochrome *c* oxidase subunit 1 (COI) full database of barcoding sequences, we found a 99.68% match in sequence similarity to a specimen identified as *Austrolimnophila* spp. (specimen ID MPG2107-22) sequenced at the Centre for Biodiversity Genomics at Guelph, Ontario CA.

### Infrared imaging and temperature measurements

We assessed the cold tolerance ability of snow flies (n = 77 flies) and other crane flies (n = 16 flies) collected in by placing them on a TECA Model AHP-301CPV cold plate preset to cool at a rate of 0.80 °C/min for 25 minutes, after which the temperature increased to 2 °C. Each trial took place in a cold room at an ambient temperature of 5 °C. To prevent escape, all flies were held within an aluminum ring coated with Rain-X or liquid graphite lubricant. On occasion, a paint brush or canned air was used to keep the fly within the ring. Each trial was recorded using a FLIR T860 infrared camera (30 FPS) elevated 22 cm from the surface of the cold plate. The object and atmosphere parameters were adjusted from factory settings to account for the distance of the lens to the surface of the cold plate, the reflected temperature of the cold plate, the atmospheric temperature of the cold room, and the relative humidity of the cold room. This was necessary to ensure accuracy of temperature readings. To validate the accuracy of the thermal camera, we attached a Physitemp MT-29/1HT needle microprobe to 11 crane flies using UV glue or Vaseline and recorded their surface temperature using an Onset HOBO UX120-014M 4-Channel thermocouple data logger throughout the trial (**Fig. S1**). After each trial was complete, the supercooling point (SCP) was determined using the FLIR ResearchIR program with a 1×1 pixel ROI centered on the abdomen of each fly. The supercooling point was indicated by a rapid temperature increase of at least 1 °C within the fly’s body. Visually, the fly was observed to rapidly change color when monitoring the video in real time or using the ResearchIR software, as indicated in **Fig. 2B, E**. The survival of the fly and number of legs lost was recorded after the trial. In **Fig. 2B**, we determined the temperature of a representative snow fly (SF0181; **Table S1**) in a downsampled thermal video (0.5 fps) by finding the minimum temperature of the abdomen within a 3×3 pixel ROI based on its 2D position. Note that the temperature values were extracted from frame-specific temperature maps, which we downloaded from the FLIR ResearchIR software. In **Fig. 2E**, we determined the temperature of another snow fly (SF0166; **Table S1**) in a non-downsampled video (30 fps) by recording the minimum temperature of the abdomen (frame-by-frame) using a 3×3 pixel ROI in Research IR.

### Inducing autotomy by mechanical stimulation of the leg

To provoke leg autotomy, we briefly anesthetized each fly using CO_2_ and gently grasped the tarsal segment of one leg using pean forceps. Anesthetization was necessary to slow the fly and ensure accurate capture of the leg by forceps. Based upon previous observations that all limbs are capable of detachment, we grasped any available limb for the experiment. When the fly had regained locomotor ability after anesthetization, the tibia and tarsus of the captive limb was repeatedly prodded using thumb forceps or a fine paint brush for one minute. This was repeated every five minutes for 15 minutes. Snow fly trials (n = 8 flies) took place in the same cold room used for thermal imaging (5 °C). Crane fly trials (n = 13 flies) took place outside of the cold room (21 °C). Additionally crane fly wings were clipped to prevent flight. Some flies autotomized their legs during the initial grasping of the leg with forceps. These were recorded as instances of autotomy. Afterward, we conducted cold tolerance experiments using the same flies to see whether freezing initiated autotomy. Two snow flies had died between experiments, reducing the number of flies used for the cold tolerance experiments (n = 6).

### Motion tracking and quantification of locomotion

To determine body velocity and trajectory of the snow fly in **Fig. 2B & C**, we manually tracked the abdomen of a representative snow fly (SF0181, **Table S1**) in a thermal video that were downsampled from 30 fps to 0.5 fps. We converted the position of the abdomen in each frame from pixels to millimeters using a conversion factor based on the measured diameter of the aluminum ring in pixels and millimeters (600 pixels and 70 millimeters, respectively). Instantaneous body velocity was calculated in python using the following equation:

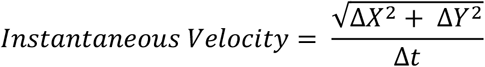

Where Δ*X* and Δ*Y* are the frame-by-frame change in the 2D position of the fly and Δ*t* is the time interval between frames.

Locomotor ability of snow flies and other crane flies was assessed by examining whether snow flies (n = 17 flies) and crane flies (n = 5 flies) engaged in coordinated movement at 4 °C cold-plate temperature intervals during a 25-minute imaging session. The temperature was assessed using a 3×3 ROI cursor fixed on the cold plate. We manually scored fly locomotion within 6 different windows (cold plate temperatures of 3, -1, -9, -13, -17 +/- 0.5 °C).

### Confocal imaging of intact and detached legs

Intact and autotomized legs were mounted in VECTASHIELD media (Vector Laboratories) and imaged with a 20x objective on a Confocal Olympus FV1000 using a far-red laser (633nm) to collect z-stacks of cuticle auto-fluorescence. We processed maximum-projection images in FIJI (Schindelin et al. 2012).

### Statistical analysis

We used scripts written in Python to perform analyses used for **Figs. 2, 3**. We tested for statistically significant differences between the two groups using two sample t-tests (**Figs. 2, 3**), performed using Python and Google Sheets.

### Analysis code availability

Python code used to determine snow fly temperature, body velocity, and trajectory, as well as, for generating plots is located at https://github.com/Prattbuw/CODE-Snow-flies-self-amputate-freezing-limbs-to-sustain-behavior-at-sub-zero-temperatures.

## Supplemental Information

**Fig. S1.**
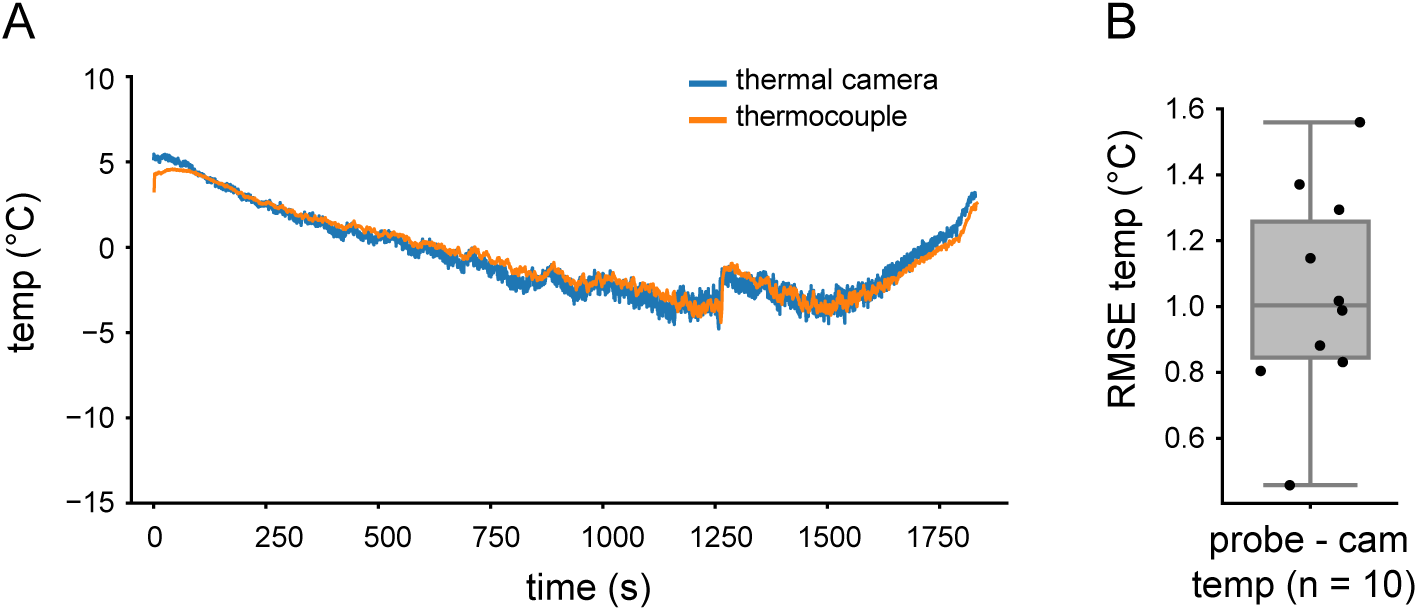
Temperatures recorded using thermal imaging are similar to temperatures recorded using a thermocouple. (**A**) Example of thermal imaging recorded temperature vs thermocouple temperature for a winged crane fly during a cold tolerance experiment. (**B**) The root mean squared error (RMSE) between the fly’s temperature recorded by the FLIR camera and that recorded by a thermocouple probe for 10 trials (each black dot corresponds to the RMSE of a trial). The probe recorded the fly’s temperature on average about 1°C higher than the FLIR camera.

**Video S1**: Examples of snow fly locomotion in the wild. We have observed snow flies walking on the snow at temperatures close to -10 °C.

**Video S2**: Example of thermal imaging of a walking snow fly in the wild (on snow).

**Video S3**: Snow fly vs. crane fly activity at -5 °C. Crane fly locomotive ability diminished when exposed to the ambient cold room temperature of 5 °C. Almost all locomotion ceased at temperatures below 3 °C on the cold plate.

**Video S4**: Example experimental trial, demonstrating the SCP. Snow flies did not enter a state of paralysis due to cold exposure, allowing them to sustain locomotion until the moment that they freeze. Freezing is indicated by a rapid change in color that corresponds to an increase in body temperature.

**Video S5**: Snow fly autotomy is triggered by freezing of the leg. Autotomy of the freezing leg occurs within seconds and can occur before the freezing hemolymph, initiated in the tarsus, reaches the femur-trochanter joint. As failure of leg detachment can lead to death, some flies autotomize more than one freezing leg to prolong their survival or autotomize the leg while in motion.

**Video S6**: Extreme example of a snow fly in the wild missing legs.

**Video S7**: Crane fly autotomy is triggered by mechanical stimuli while snow fly autotomy is triggered by freezing stimuli. Mechanical autotomy experiments were conducted prior to cold tolerance experiments.

**Table S1**: Snow fly collection location and species data sheet.

**Table S2**: Crane fly collection location and species data sheet.

