## Supplementary material for "Snow flies self-amputate freezing limbs to sustain behavior at sub-zero temperatures": table s1

**Table S1: Snow fly collection location and species data sheet.**

| fly_id | sex | species (DNA ID) | collection_date | latitude | longitude | elevation (m) |
|---|---|---|---|---|---|---|
| SF0056 | female | alexandriana | 12/29/20 | 47.439167 | -120.37222 | 1524 |
| SF0070 | male | alexandriana | 12/29/20 | 47.439167 | -120.37222 | 1524 |
| SF0075 | male | alexandriana | 12/29/20 | 47.439167 | -120.37222 | 1524 |
| SF0077 | male | excavata | 12/29/20 | 47.439167 | -120.37222 | 1524 |
| SF0079 | male | alexandriana | 12/29/20 | 47.439167 | -120.37222 | 1524 |
| SF0080 | male | alexandriana | 12/29/20 | 47.439167 | -120.37222 | 1524 |
| SF0063 | male | alexandriana | 12/31/20 | 46.930278 | -121.4525 | 1889.76 |
| SF0066 | female | alexandriana | 12/31/20 | 46.930278 | -121.4525 | 1889.76 |
| SF0085 | female | alexandriana | 1/9/21 | 48.618056 | -120.32556 | 1940.357 |
| SF0087 | female | alexandriana | 1/9/21 | 48.618056 | -120.32556 | 1940.357 |
| SF0130 | male | excavata | 11/7/2021 | 47.37664 | -121.45107 | 1182.624 |
| SF0131 | male | alexandriana | 11/7/2021 | 47.37664 | -121.45107 | 1182.624 |
| SF0133 | female | alexandriana | 11/7/2021 | 47.37664 | -121.45107 | 1182.624 |
| SF0134 | male | alexandriana | 11/7/2021 | 47.37664 | -121.45107 | 1182.624 |
| SF0129 | male | alexandriana | 11/7/2021 | 47.37664 | -121.45107 | 1182.624 |
| SF0135 | female | alexandriana | 11/7/2021 | 47.37664 | -121.45107 | 1182.624 |
| SF0132 | male | alexandriana | 11/7/2021 | 47.37664 | -121.45107 | 1182.624 |
| SF0140 | female | alexandriana | 11/21/2021 | 46.79483 | -121.71869 | 1859.28 |
| SF0141 | female | alexandriana | 11/21/2021 | 46.79483 | -121.71869 | 1859.28 |
| SF0142 | male | alexandriana | 11/21/2021 | 46.79483 | -121.71869 | 1859.28 |
| SF0145 | male | alexandriana | 11/21/2021 | 46.79483 | -121.71869 | 1859.28 |
| SF0146 | male | alexandriana | 11/21/2021 | 46.79483 | -121.71869 | 1859.28 |
| SF0147 | male | alexandriana | 11/21/2021 | 46.79483 | -121.71869 | 1859.28 |
| SF0148 | male | alexandriana | 11/21/2021 | 46.79483 | -121.71869 | 1859.28 |
| SF0149 | female | excavata | 11/21/2021 | 46.79483 | -121.71869 | 1859.28 |
| SF0150 | female | alexandriana | 11/24/2021 | 48.84931 | -121.69953 | 1676.4 |
| SF0151 | female | excavata | 11/24/2021 | 48.84931 | -121.69953 | 1676.4 |
| SF0153 | male | excavata | 11/24/2021 | 48.84931 | -121.69953 | 1676.4 |
| SF0154 | female | alexandriana | 11/24/2021 | 48.84931 | -121.69953 | 1676.4 |
| SF0156 | male | alexandriana | 12/5/2021 | 47.3776 | -121.34073 | 1554.48 |
| SF0157 | male | alexandriana | 12/5/2021 | 47.3776 | -121.34073 | 1554.48 |
| SF0158 | male | alexandriana | 12/5/2021 | 47.3776 | -121.34073 | 1554.48 |

| | | | | | | |
|---|---|---|---|---|---|---|
| SF0160 | female | alexandriana | 12/12/2021 | 47.3989 | -121.42386 | 1158.24 |
| SF0163 | female | excavata | 12/12/2021 | 47.3989 | -121.42386 | 1158.24 |
| SF0164 | female | alexandriana | 12/12/2021 | 47.3989 | -121.42386 | 1158.24 |
| SF0165 | male | excavata | 12/12/2021 | 47.3989 | -121.42386 | 1158.24 |
| SF0166 | female | macnabeana | 12/12/2021 | 47.3989 | -121.42386 | 1158.24 |
| SF0168 | female | excavata | 12/12/2021 | 48.84931 | -121.69953 | 1676.4 |
| SF0169 | female | alexandriana | 12/12/2021 | 48.84931 | -121.69953 | 1676.4 |
| SF0170 | male | excavata | 12/12/2021 | 48.84931 | -121.69953 | 1676.4 |
| SF0171 | female | excavata | 12/8/2021 | 48.84931 | -121.69953 | 1676.4 |
| SF0173 | male | alexandriana | 12/22/2021 | 47.7444 | -121.02939 | 1687.068 |
| SF0174 | male | macnabeana | 12/22/2021 | 47.7444 | -121.02939 | 1687.068 |
| SF0175 | male | alexandriana | 12/22/2021 | 47.7444 | -121.02939 | 1687.068 |
| SF0176 | male | alexandriana | 12/22/2021 | 47.7444 | -121.02939 | 1687.068 |
| SF0177 | female | macnabeana | 12/22/2021 | 47.7444 | -121.02939 | 1687.068 |
| SF0178 | female | albertensis | 12/22/2021 | 47.7444 | -121.02939 | 1687.068 |
| SF0179 | male | alexandriana | 12/22/2021 | 47.7444 | -121.02939 | 1687.068 |
| SF0180 | female | alexandriana | 12/22/2021 | 47.7444 | -121.02939 | 1687.068 |
| SF0181 | female | alexandriana | 12/22/2021 | 47.7444 | -121.02939 | 1687.068 |
| SF0182 | female | alexandriana | 12/22/2021 | 47.7444 | -121.02939 | 1687.068 |
| SF0183 | male | albertensis | 12/22/2021 | 47.7444 | -121.02939 | 1687.068 |
| SF0184 | female | alexandriana | 12/22/2021 | 47.7444 | -121.02939 | 1687.068 |
| SF0185 | female | alexandriana | 12/22/2021 | 47.7444 | -121.02939 | 1687.068 |
| SF0186 | male | alexandriana | 12/22/2021 | 47.7444 | -121.02939 | 1687.068 |
| SF0187 | male | albertensis | 12/22/2021 | 47.7444 | -121.02939 | 1687.068 |
| SF0191 | female | alexandriana | 1/5/2022 | 47.44829 | -121.4279 | 1014.984 |
| SF0192 | female | alexandriana | 1/5/2022 | 47.44829 | -121.4279 | 1014.984 |
| SF0193 | male | alexandriana | 1/5/2022 | 47.44829 | -121.4279 | 1014.984 |
| SF0194 | female | alexandriana | 1/5/2022 | 47.44829 | -121.4279 | 1014.984 |
| SF0195 | male | alexandriana | 1/5/2022 | 47.44829 | -121.4279 | 1014.984 |
| SF0196 | female | macnabeana | 1/5/2022 | 47.44829 | -121.4279 | 1014.984 |
| SF0197 | female | alexandriana | 1/5/2022 | 47.44829 | -121.4279 | 1014.984 |
| SF0198 | male | alexandriana | 1/5/2022 | 47.44829 | -121.4279 | 1014.984 |
| SF0200 | male | albertensis | 1/5/2022 | 47.44829 | -121.4279 | 1014.984 |

| | | | | | | |
|---|---|---|---|---|---|---|
| SF0201 | female | alexandriana | 1/5/2022 | 47.44829 | -121.4279 | 1014.984 |
| SF0202 | female | alexandriana | 1/5/2022 | 47.44829 | -121.4279 | 1014.984 |
| SF0203 | female | excavata | 1/5/2022 | 47.44829 | -121.4279 | 1014.984 |
| SF0204 | male | alexandriana | 1/5/2022 | 47.44829 | -121.4279 | 1014.984 |
| SF0206 | male | excavata | 1/14/2022 | 48.853556 | -121.69314 | 1290.523 |
| SF0207 | female | excavata | 1/14/2022 | 48.853556 | -121.69314 | 1290.523 |
| SF0217 | female | alexandriana | 1/16/2022 | 48.71551 | -121.99175 | 1687.373 |
| SF0220 | male | alexandriana | 1/30/2022 | 47.44829 | -121.4279 | 1014.984 |
| SF0221 | female | excavata | 1/30/2022 | 47.44829 | -121.4279 | 1014.984 |
| SF0222 | male | alexandriana | 1/30/2022 | 47.44829 | -121.4279 | 1014.984 |
| SF0244 | female | alexandriana | 4/6/2022 | 47.458611 | -121.41733 | 1866.9 |
| SF0245 | male | alexandriana | 4/10/2022 | 47.456694 | -121.41464 | 1758.086 |
