## Supplementary material for "Snow flies self-amputate freezing limbs to sustain behavior at sub-zero temperatures": table s2

**Table S2: Crane fly collection location and species data sheet.**

| fly_id | sex | species (DNA ID) | collection_date | latitude | longitude | elevation (ft) |
|---|---|---|---|---|---|---|
| CF0005 | male | Austrolimnophila | 5/29/2022 | 47.33088 | -122.6372 | 196 |
| CF0006 | female | Austrolimnophila | 5/29/2022 | 47.33088 | -122.6372 | 196 |
| CF0008 | male | Austrolimnophila | 5/29/2022 | 47.33088 | -122.6372 | 196 |
| CF0009 | male | Austrolimnophila | 5/29/2022 | 47.33088 | -122.6372 | 196 |
| CF0010 | male | Austrolimnophila | 5/29/2022 | 47.33088 | -122.6372 | 196 |
| CF0022 | male | | 7/18/2022 | 47.33088 | -122.6372 | 196 |
| CF0024 | female | | 7/20/2022 | 47.33088 | -122.6372 | 196 |
| CF0025 | male | | 7/20/2022 | 47.33088 | -122.6372 | 196 |
| CF0029 | female | | 8/1/2022 | 47.33088 | -122.6372 | 196 |
| CF0032 | female | | 8/7/2022 | 47.68497 | -122.2981 | 334 |
| CF0033 | female | | 8/25/2022 | 47.69204 | -122.2828 | 285 |
| CF0035 | male | | 8/28/2022 | 46.458704 | -122.7665 | 139 |
| CF0036 | female | | 8/28/2022 | 46.458704 | -122.7665 | 139 |
| CF0038 | male | | 8/30/2022 | 47.69204 | -122.2828 | 285 |
| CF0039 | male | | 8/30/2022 | 47.69204 | -122.2828 | 285 |
| CF0040 | female | | 8/30/2022 | 47.69204 | -122.2828 | 285 |
